## Supporting Info Tables and Figures for "Information Decay and Enzymatic Information Recovery for DNA Data Storage"

**Supporting Information 1: Sets of DNA used for this work.**

For all of the following repair reactions, the following procedure was followed: Individual components were added on ice to make master mix: (ThermoPol buffer, dNTP, NAD+, enzyme 1, enzyme 2, enzyme n (ratio: 10:4:2:x:y:z). For repair reactions, 18.8 µL mili-Q water, 6.25 µL DNA (1 ng/µL), and 1.57 µL master mix were added (for control samples, the 1.57 µL master mix were exchanged for water). After the addition of all components, reaction vials were shaken by hand and centrifuged. Repair was carried out in 30 °C incubator for 15 min. Samples were subsequently centrifuged again. PCR quantification was performed by diluting 1 µL of incubated DNA in 499 µL water.

***Supplementary Table 1:*** *DNA used for this work. Shown are properties of each set of DNA*

| **DNA name** | **Encoded information** | **Size of file [bytes]** | **Size of file [unique DNA strands]** | **Length of DNA strands** |
|---|---|---|---|---|
| **Model DNA** | - | - | 1 | 113 |
| **File 1 DNA** | Jpg image | 115,394 | 7,373 | 150 |

**Supporting Information 2: Selection of enzymes investigated for repair.**

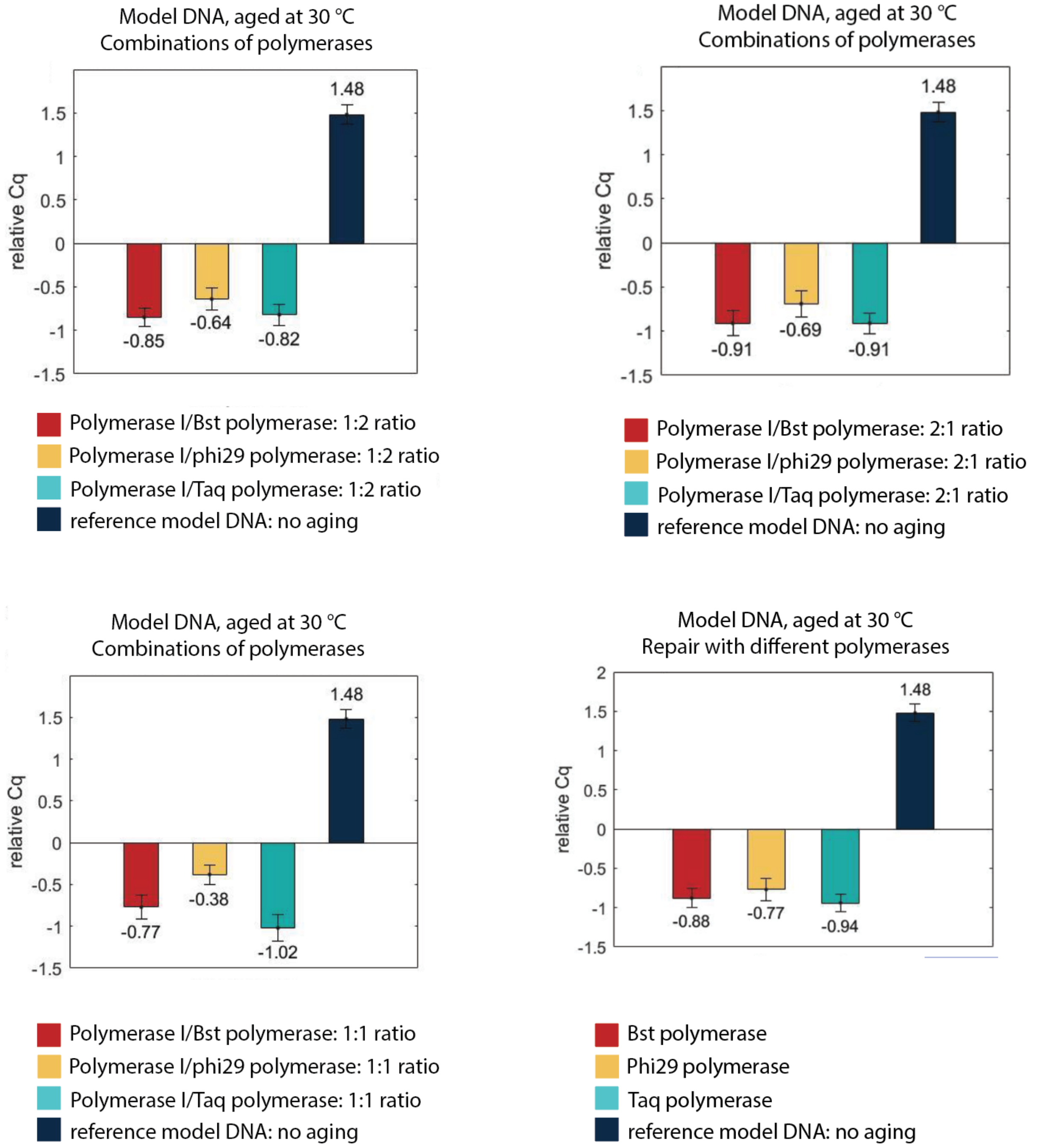


***Supplementary Figure 1: The effect of polymerases on repair was tested.*** *A repair reaction consisted of an endonuclease (no APE 1), a polymerase and a ligase. Different polymerase combinations were investigated but did not improve repair. This shows that in the absence of APE1, no matter what polymerase is chosen, no repair takes place. Relative cq on the y-axis represents the relative number of cycles to the aged (and not repaired) sample.*

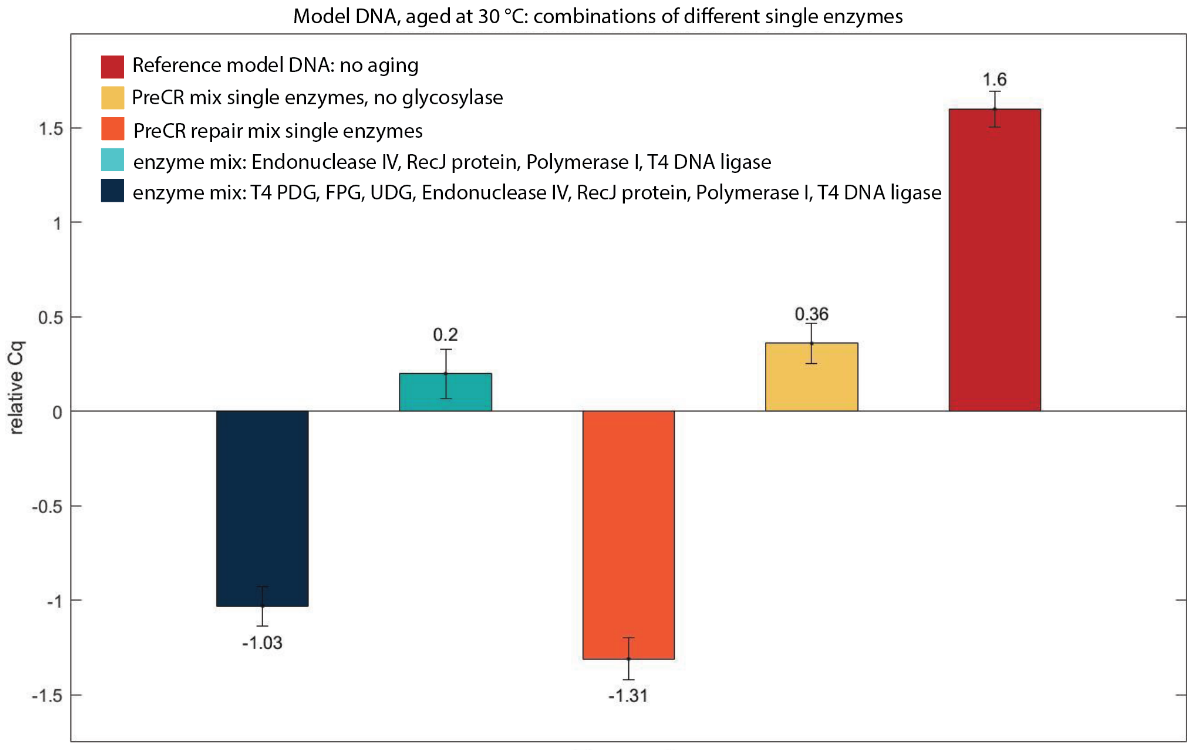


***Supplementary Figure 2: Enzyme mix repair tests.*** *Lindahl enzymes for the BER pathway as well as enzymes contained in PreCR® repair mix tested as a combination of single enzymes. Relative cq on the y-axis represents the relative number of cycles to the aged (and not repaired) sample of model DNA.*

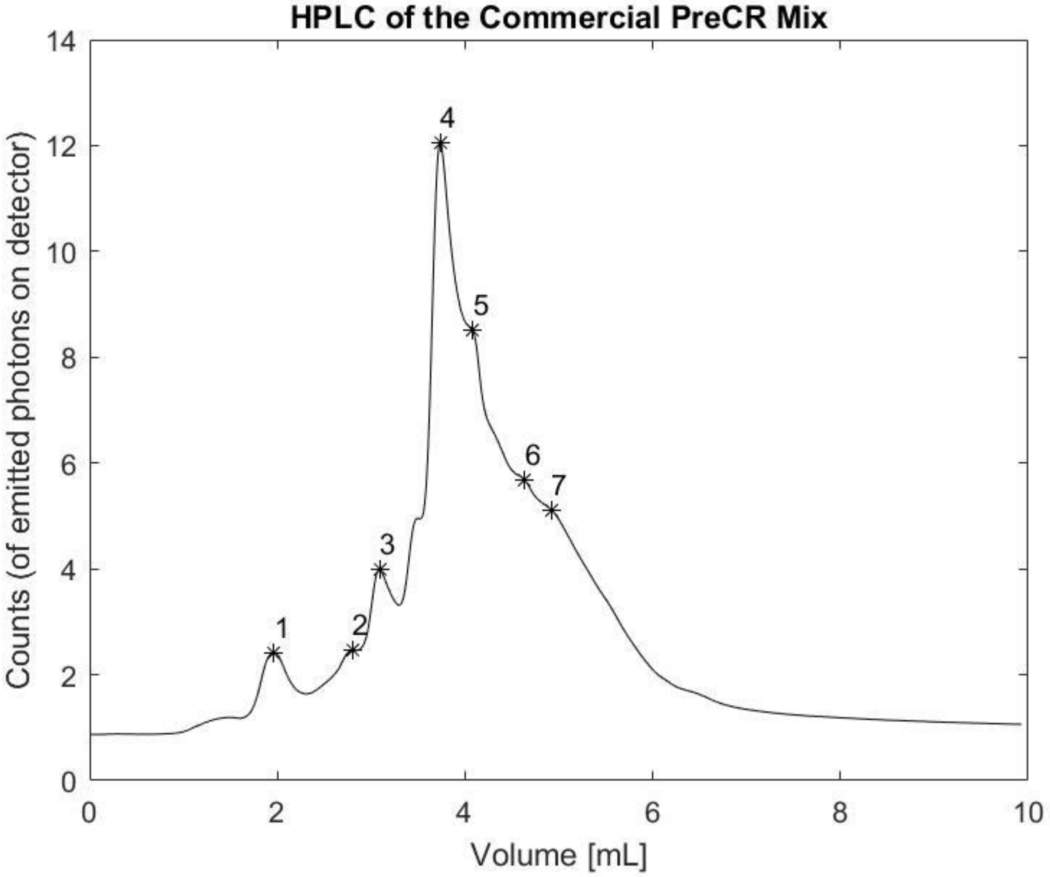


***Supplementary Figure 3: HPLC measurement of PreCR® repair mix.*** *20 µL of the mix were analyzed and seven peaks corresponding to 7 enzymes in PreCR® repair mix were observed. Enzymes appear in the order: 1) Bst polymerase 2) Taq ligase 3) FPG glycosylase 4) Endonuclease IV 5) Endonuclease VIII 6) UDG glycosylase 7) T4 PDG glycosylase. The peaks were converted into relative volumes of enzyme present (as shown in Supplementary Table D.2)*

***Supplementary Table 2: Composition of PreCR® repair mix.*** *Calculated relative amounts of single enzymes in PreCR repair mix based on HPLC results (Supplementary Figure D.3)*

| **Peak** | **Enzyme** | **Relative amount in PreCR mix** | **Number of tryptophan molecules** | **Molecular weight kDa** |
|---|---|---|---|---|
| 1 | Bst Polymerase | 1 | 5 | 99 |
| 2 | Taq Ligase | 1 | 5 | 77 |
| 3 | FPG (glycosylase) | 2 | 4 | 43 |
| 4 | Endonuclease IV | 6 | 4 | 31 |
| 5 | Endonuclease VIII | 4 | 4 | 30 |
| 6 | UDG (glycosylase) | 2 | 6 | 26 |
| 7 | T4 PDG (glycosylase) | 10 | 1 | 16 |

***Supplementary Table 3: Enzyme mixes tested for repair.*** *Different volume ratios of repair mixes containing single enzymes tested.*

| **Enzymes** | ***DNA*** | ***Relative cq*** |
|---|---|---|
| polymerase I, T4 DNA ligase, RecJ protein (ratio: 1:2:4) | *File 1, aged at 25 °C* | *-0.64 ± 0.04* |
| polymerase I, T4 DNA ligase, RecJ protein (ratio: 2:1:4) | *File 1, aged at 25 °C* | *-0.77 ± 0.07* |
| Endonuclease VIII, APE1, Bst polymerase, Taq ligase, T4 polynucleotide kinase | *File 1, aged at 30 °C* | *+0.80 ± 0.04* |
| Endonuclease IV, APE1, Bst polymerase, Taq ligase, T4 polynucleotide kinase | *File 1, aged at 30 °C* | *+1.24 ± 0.10* |
| Endonuclease IV, Endonuclease VIII, Bst polymerase, Taq ligase, T4 polynucleotide kinase | *File 1, aged at 30 °C* | *+1.17 ± 0.06* |
| Endonuclease IV, Endonuclease VIII, APE1, Taq ligase, T4 polynucleotide kinase | *File 1, aged at 30 °C* | *+0.35 ± 0.12* |
| Endonuclease IV, Endonuclease VIII, APE1, Bst Polymerase, T4 polynucleotide kinase | *File 1, aged at 30 °C* | *+1.28 ± 0.08* |
| Endonuclease IV, Endonuclease VIII, APE1, Bst Polymerase, Taq Ligase | *File 1, aged at 30 °C* | *+1.16 ± 0.08* |
| Endonuclease IV, Endonuclease VIII, APE1, Bst Polymerase, Taq Ligase, T4 polynucleotide kinase | *File 1, aged at 30 °C* | *+1.12 ± 0.11* |
| APE1, Bst Polymerase, T4 DNA ligase | *File 1, aged at 30 °C* | *+1.00 ± 0.17* |

**Supporting Information 3: Fragmentation size and melting point analysis.**

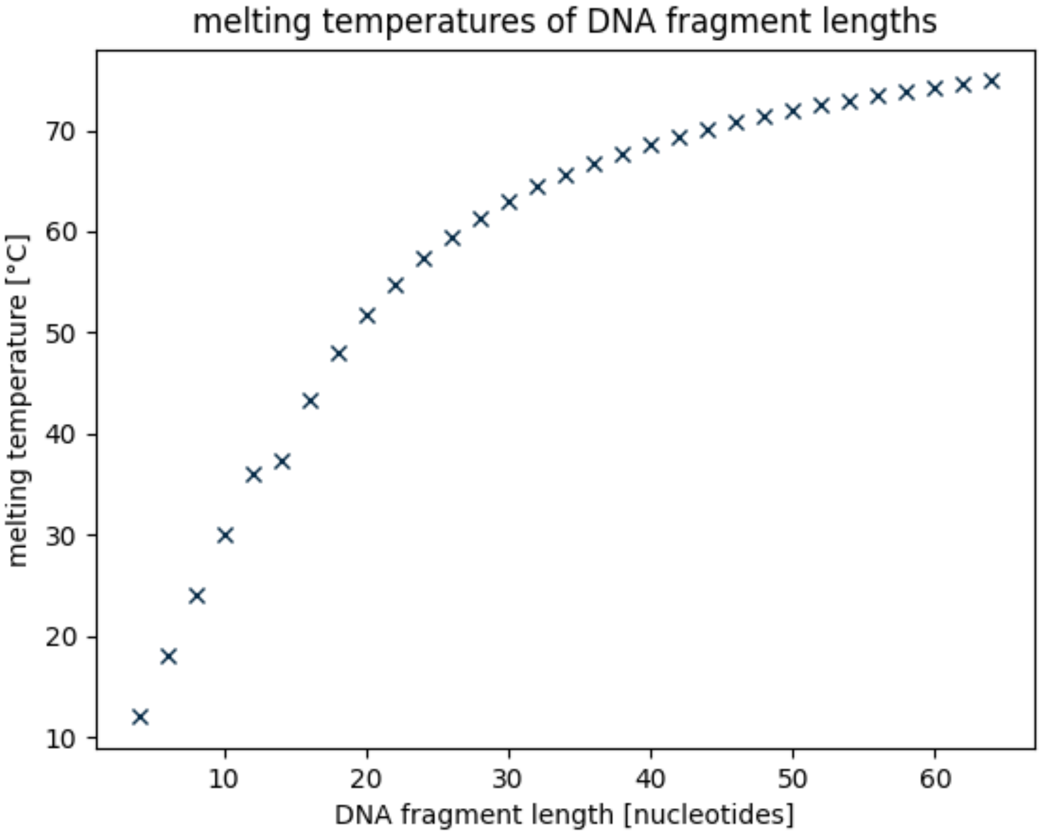


***Supplementary Figure 4: Melting temperatures of DNA fragments.*** *Basic melting temperatures of different DNA fragment lengths with 50% GC content. Calculations were based on the analysis tools by Kibbe et al.*[228]

As the length of DNA fragments determines the respective melting point of the fragment, the position of nicks may affect the fragmentation analysis, too. This is, because the shorter the fragment after nicking, the lower the melting temperature of that fragment. If aging proceeds at 30 °C (as was in our experiments), fragments shorter than 10 nucleotides will denature during aging procedures already. Consequentially, repair of the denatured fragment will no longer be possible, as repair can only proceed at nicks in double stranded DNA. This explains the importance of choosing aging temperatures according to the fragment length that is to be analyzed. If aging were to be carried out at 70 °C (i.e., in enhanced aging experiments), all fragments shorter than 40 nucleotides would dissociate and in turn, repair of these samples could no longer take place.

Returning to our analysis at 30 °C: What would happen, if a DNA strand is nicked more than once? Assuming the original sequence length is around 150 nucleotides (as was the case in our experiments), two evenly distributed nicks would render three DNA fragments with a length of 50 nucleotides. However, as we increase the number of nicks (evenly distributed within our DNA strand for simplicity reasons), the resulting fragments become smaller with every additional nick. At 14 nicks, each fragment is only 10 nucleotides long and would dissociate during aging. Besides aging, another bottleneck step is repair itself, which proceeds at 37 °C. At this temperature, all fragments smaller than 15 nucleotides would dissociate before repair can begin. These temperature considerations prove especially relevant for repair-related reactions and illustrate why analysis of short fragments can potentially not be quantitatively accurate.

### Supporting Information 4: Simulation details

Simulations (Figure 5) were performed using the python script below. The major assumptions are that the DNA synthesis bias (= number distribution of copies of unique oligos in the pool) follows a normal distribtion with a standard deviation equal to 0.32 times the physical redundancy, as described in Chen. et al. *Nat. Commun.* **11**, 3246 (2020).

The PCR bias was modelled assuming that the per-strand PCR efficiency is follows a normal distribution, with an average of 1.85 and a standard deviation of 0.07, as described in Heckel et al. *Sci. Rep*. **9**, 9663 (2019).

```
# Simulating DNA Damage and Repair

import numpy as np
import pandas as pd

from multiprocessing import Pool

rng = np.random.default_rng()

# simulation settings
num_seq = 10000                 # number of unique sequences
phys_red_max = 50               # upper limit of physical redundancy to simulate
r_vals = [0.90, 0.50, 0.10, 0.0]       # repair ratios to simulate
d_vals = [0.9375, 0.875, 0.75, 0.50]   # damage ratios to simulate
iterations = 1000               # number of simulations per unique combination of parameters
def simulate(num_seq, phys_red, dam_fac, rep_fac):

    #
    # SYNTHESIS
    #
    # Find the actual physical redundancy based on a normal distribution
    pr = rng.normal(loc = phys_red, scale = 0.32*phys_red, size = num_seq)
    pr = np.rint(pr)
    pr[pr < 0] = 0

    #
    # PCR
    #
    # simulate the change in physical redundancy caused by PCR bias
    num_cyc = 30
    eff_PCR = rng.normal(loc = 1.85, scale = 0.07, size = num_seq)
    bias_PCR = eff_PCR/np.mean(eff_PCR)
    pr_pcr = pr * (bias_PCR ** num_cyc)
    pr_pcr = np.rint(pr_pcr).astype('int')
    pr[pr < 0] = 0

    # re-normalize to desired mean physical redundancy (== dilution)
    pr_pcr_dil = rng.binomial(pr_pcr, phys_red/np.mean(pr_pcr))

    #
    # DAMAGING
    #

    # get number of damaged oligos per sequence
    pr_damaged = rng.binomial(pr_pcr_dil, dam_fac)
    pr_post_damage = pr_pcr_dil - pr_damaged
    lost_seqs_post_damage = np.count_nonzero(pr_post_damage == 0)

    #
    # REPAIR
    #
    pr_repaired = rng.binomial(pr_damaged, rep_fac)
    pr_post_repair = pr_post_damage + pr_repaired
    lost_seqs_post_repair = np.count_nonzero(pr_post_repair == 0)
```

```
    return lost_seqs_post_damage, lost_seqs_post_repair

def process_simulation(rep_fac):
    for dam_fac in d_vals:
        for phys_red in range(1, phys_red_max + 1):
            list_num_seq = np.zeros(iterations)
            list_phys_red = np.zeros(iterations)
            list_dam_fac = np.zeros(iterations)
            list_rep_fac = np.zeros(iterations)
            list_lost_seqs_damage = np.zeros(iterations)
            list_lost_seqs_repair = np.zeros(iterations)

            for iter in range(iterations):
                lost_seqs_post_damage, lost_seqs_post_repair = simulate(num_seq, phys_red, dam_fac, rep_fac)
                list_num_seq[iter] = num_seq
                list_phys_red[iter] = phys_red
                list_dam_fac[iter] = dam_fac
                list_rep_fac[iter] = rep_fac
                list_lost_seqs_damage[iter] = lost_seqs_post_damage
                list_lost_seqs_repair[iter] = lost_seqs_post_repair

            # collect data from all iterations and write to file
            df2 = pd.DataFrame({
                'num_seq': list_num_seq,
                'phys_red': list_phys_red,
                'dam_fac': list_dam_fac,
                'rep_fac': list_rep_fac,
                'lost_useq_post_damage': list_lost_seqs_damage,
                'lost_useq_post_repair': list_lost_seqs_repair,
            })
            df2.to_csv(f'files/nonidpcr_{str(num_seq/1000)}k_{str(rep_fac)}.csv', mode='a', header=False)
if __name__ == '__main__':
    pool = Pool()
    pool.map(process_simulation, r_vals)
```
